# The evolution of wasp mimicry and biogeography in the genus *Temnostoma* (Diptera: Syrphidae)

**DOI:** 10.1101/2024.08.27.609869

**Authors:** Jiří Hadrava, Jan Klečka, Kevin Moran, Irena Klečková, Scott Kelso, Claudia Etzbauer, Jeffrey H. Skevington, Ximo Mengual

## Abstract

One of the most remarkable examples of Batesian mimicry occurs in the genus *Temnostoma* Le Peletier & Audinet-Serville, 1828 (Diptera: Syrphidae). Adults of this genus have an overall resemblance to hymenopterans combined with behavioural mimicry (they move the fore legs in front of the head mimicking hymenopteran antennae). While some species of *Temnostoma* are considered perfect mimics of social wasps, other species have a darker colour pattern and are rather imperfect mimics. Both colour morphs are widespread through the Holarctic. Here, we infer for the first time the evolutionary history of the genus with special focus on the evolution of mimicry and biogeography.

With material covering 75 % of known species of *Temnostoma* and both colour morphs from each biogeographical region, we inferred a molecular phylogeny based on six molecular markers (mitochondrial protein-coding COI gene, nuclear 28s rRNA gene, and four nuclear protein-coding genes: AATS, CK1, TULP, and RBP-15). Using Bayesian inference, we obtained a highly resolved phylogenetic tree supporting the monophyly of the genus *Temnostoma* as a sister group of genus *Takaomyia* Hervé-Bazin, 1914. Within *Temnostoma*, *Te. daochus* and *Te. barberi* (two Nearctic species with strikingly different mimicry patterns) were found to be closely related to each other and together form a lineage sister to the rest of the genus.

Our results suggest that the behavioural mimicry of wasp antennae is a plesiomorphic state inherited from a common ancestor that includes the genera *Temnostoma* and *Takaomyia*. Within *Temnostoma*, the dark colour pattern (imperfect mimicry) appeared to be an ancestral state and perfect wasp mimicry evolved two times independently within the genus. In some species inhabiting northern parts of the Holarctic, secondary darkening and consequent degradation of the wasp mimicry appeared. This indicates high evolutionary plasticity and ongoing selection pressure on morphological characters related to mimicry in hover flies.

## 1. Introduction

With more than 6200 described species (Skevington et al. 2019), hover flies (Diptera: Syrphidae) are one of the largest and best-known families of Diptera. They inhabit a wide range of habitats across all continents excluding Antarctica. Hover flies are an important group of pollinators (Ssymank et al. 2008, Klečka et al. 2018) and bioindicators (Sommaggio 1999, Vujić et al. 2016). However, they are best known for their Batesian mimicry of aculeate Hymenoptera (Waldbauer 1970, Dlusski 1984, Drees 1997). To avoid predation pressure caused by birds or other visually oriented predators, many species of hover flies exhibit contrasting colour patterns, morphological, or even behavioural adaptations that make them resemble various groups of bees or wasps (Waldbauer 1970, Dlusski 1984, Drees 1997). One of the most notable examples of Batesian mimicry can be found in the genus *Temnostoma* Le Peletier & Audinet-Serville, 1828.

*Temnostoma* has been traditionally classified in the subfamily Eristalinae, tribe Milesiini, and subtribe Temnostomina. Still, the higher systematics of Syrphidae has been questioned several times in recent years (e.g. Young et al. 2016, Pauli et al. 2018, Moran et al. 2022). Based on the most recent phylogeny of Syrphidae (Moran et al. 2022), *Temnostoma* is placed within the subfamily Eristalinae, but the tribe Milesiini as well as Temnostomina are not resolved as monophyletic groups. While Milesiini is found to be a rather artificial group comprising various Eristalinae with rather large body sizes, Temnostomina is a non-natural group comprising various genera exhibiting behavioural mimicry of Hymenopteran antennae. The only other genus of Temnostomina that appears to be truly related to *Temnostoma* in the work of Moran et al. (2022) is the East Asian genus *Takaomyia* Hervé-Bazin, 1914. The main morphological difference between genera *Temnostoma* and *Takaomyia* is the body shape, with *Takaomyia* having a narrowed, petiolate abdominal tergite 2; otherwise, the genera are very similar to each other.

Apart from wasp-like colouration, the genus *Temnostoma* is well known for its behavioural mimicry of Hymenopteran antennae using the fore legs (Waldbauer 1970, Penney et al. 2014). It is supposed that the length of antennae is an important character that bird predators are looking for when they are trying to distinguish Hymenoptera and Diptera (Bain et al. 2007). Even though *Temnostoma* exhibit wide variation in the colour of the mid and hind legs, all species of *Temnostoma* have the fore tarsomeres and apical half of fore tibiae black, and they are waved in front of the head to imitate the antennae of their Hymenopteran models (Waldbauer 1970, Penney et al. 2014). Such behaviour is generally rare in hover flies. Besides *Temnostoma*, it occurs in only a few genera (e.g. *Korinchia* Edwards, 1919, *Palumbia* Rondani, 1865, *Pterallastes* Loew, 1863, *Takaomyia* Hervé-Bazin, 1914, *Teuchocnemis* Osten-Sacken, 1876, and some species of *Spilomyia* Meigen, 1803), whose phylogenetic relationships to *Temnostoma* are poorly understood, although an unpublished hybrid enrichment phylogeny shows that these genera are in three distantly related lineages (Moran et al. in prep). Moreover, Penney et al. (2014) suggested that behavioural mimicry is typical for perfect mimics. Nevertheless, behavioural mimicry in *Temnostoma* occurs in perfect as well as in imperfect mimics, which makes them good model to study the joint evolution of morphological and behavioural mimicry.

There are 29 described species of the genus *Temnostoma*, and the genus is widespread through the Palaearctic Region, from Western Europe to East Asia, as well as in the Nearctic Region. A few Eastern Asian species extend into the Indomalayan Region.

Krivosheina (2005) divided *Temnostoma* into two subgenera: *Temnostoma (Temnostoma)* and *Temnostoma (Temnostomoides)*. *Temnostomoides* is characterised by an elongated face and wide abdominal tergites usually with two transverse yellow stripes per tergite, imitating a higher number of abdominal tergites of Hymenoptera; consequently, they have a convincing wasp look. Some *Temnostomoides* species are perfect mimics of social wasps such as *Vespula* Thomson, 1869 and *Dolichovespula* Rohwer, 1916 (Waldbauer 1970, Penney et al. 2014). Species of the subgenus *Temnostoma* have shorter face, narrower abdominal tergites, and are much darker, with at most one transverse yellow stripe per tergite. Generally, they are imperfect mimics of solitary Aculeata wasps such as Eumeninae (Shannon 1939).

The subgenus *Temnostoma (Temnostoma)* contains a single species group called *bombylans*-group (Krivosheina 2002). The subgenus *Temnostoma (Temnostomoides)* is divided into two species groups, namely the *vespiforme*-group and *apiforme*-group (Krivosheina 2004). The latter two groups can be distinguished based on characters of the male genitalia and by the scutal pattern: *vespiforme*-group species have a distinct posterolateral, triangular pale marking on each side, between the transverse suture and the scutellum, which are absent in the species belonging to the *apiforme*-group.

The subgeneric definitions by Krivosheina (2005) have some exceptions. Within *Temnostoma (Temnostomoides)*, which contains predominantly perfect wasp mimics, there are a few taxa with only a single yellow stripe per abdominal tergite, such as *Te. apiforme* (Fabricius, 1794), *Te. carens* (Gaunitz, 1936), and *Te. pallidum* Sack, 1910 (all in the *apiforme*-group). These mentioned species have a well-defined anterior stripe, but the posterior stripe is narrow and dull, sometimes reduced or almost missing. In addition, *Te. sericomyiaeforme* Portschinsky, 1886 (member of the *vespiforme*-group) always has a single yellow stripe per tergite. Nevertheless, in all the above-mentioned species the single yellow stripe is wider than typical yellow stripes of the *bombylans*-group species, making the overall appearance of the fly more prominently contrasting. Due to the northern distribution of these species, we hypothesise that this colouration could be the result of a secondary colouration reduction caused by a trade-off between mimicry and thermoregulation (Taylor et al. 2016). However, a phylogenetic framework is needed to answer whether such colouration really evolved secondarily from an ancestral wasp-like colouration (with two stripes per abdominal tergite).

Besides the above species, another species of particular interest is *Te. meridionale* Krivosheina & Mamaev, 1962. This *Temnostoma (Temnostomoides)* species resembles the *vespiforme*-group taxa based on external morphological characters such as colour pattern on thorax and abdomen; however, its male genitalia are very similar to those of the *apiforme*-group species. Consequently, Krivosheina (2005) did not assign *Te. meridionale* to any *Temnostomoides* species group. Similarly, some of the East Palaearctic and Nearctic species have not been allocated in any species group yet, and the inference of their phylogenetic affinities may help in their classification.

No phylogeny has been proposed for this genus, and thus, the relationships among species of both subgenera and the different species groups are unknown. Moreover, the biogeographic origin of the genus is unknown, together with the colonisation history of the different parts of the Holarctic Region. The presence of two distinct potential lineages (subgenera) makes *Temnostoma* a suitable model for the evolution of perfect or imperfect mimicry. Thus, the aims of our work are to infer the phylogeny of the genus *Temnostoma* based on molecular characters and to analyse the evolutionary history of wasp mimicry in this genus. First, we are interested in the ancestral morphology and the order of appearance of the various colour morphs. Second, we test the geographical origin of the genus, migrations between different parts of the Holarctic, and the occurrence of perfect and imperfect mimicry in different regions.

## 2. Material and Methods

### 2.1. Taxon sampling

We included 22 described taxa (20 species and two subspecies) and one undescribed species. For the full list of specimens, see Table 1. Our material covers the vast majority of known diversity of the subgenus *Te. (Temnostomoides)*: out of the 14 described species and two additional taxa described as subspecies, only *Te. nigrimanus* Brunetti, 1915 from India and *Te. pauperius* Speiser, 1924 from Japan are missing. In the subgenus *Te. (Temnostoma)*, we analysed eight out of the 15 described species, plus one undescribed species. The missing species are *Te. trifasciata* Robertson, 1901 from the Nearctic, and seven species from East Palaearctic and Indomalaya, namely *Te. albostriatum* Huo, Ren & Zheng, 2007 (Shaanxi Province, China), *Te. arciforma* He & Chu, 1995 (Heilongjiang and Shaanxi Provinces, China), *Te. flavidistriatum* Huo, Ren & Zheng, 2007 (Shaanxi Province, China), *Te. ningshanensis* Huo, Ren & Zheng, 2007 (Shaanxi Province, China), *Te. ravicauda* He & Chu, 1995 (Hubei Province, China), and *Te. taiwanum* Shiraki, 1930 (Taiwan). However, a comprehensive taxonomic revision is needed to reveal the real species diversity of *Te. (Temnostoma)* in the East Palaearctic and Indomalayan regions.

**Table 1.**
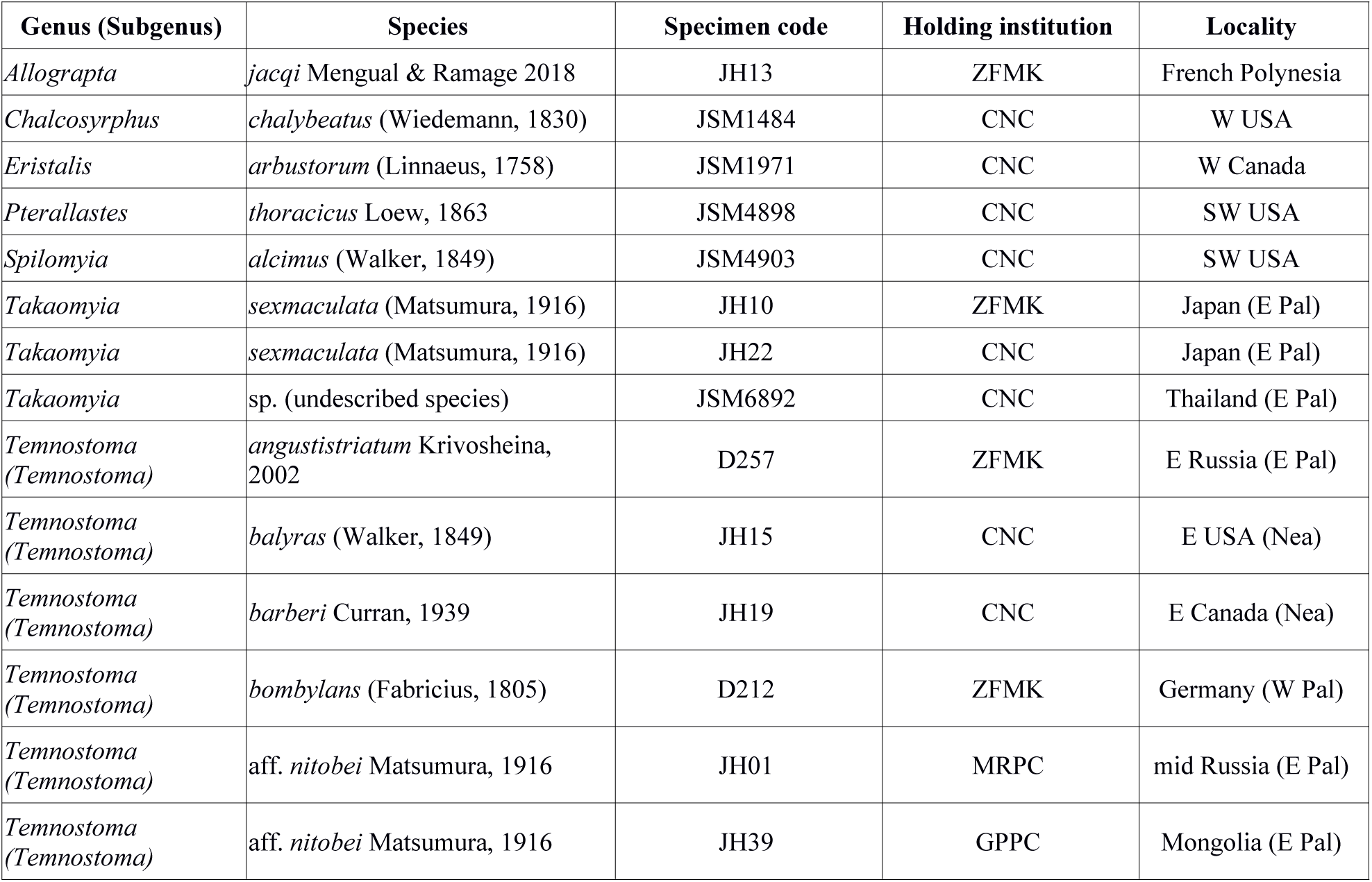
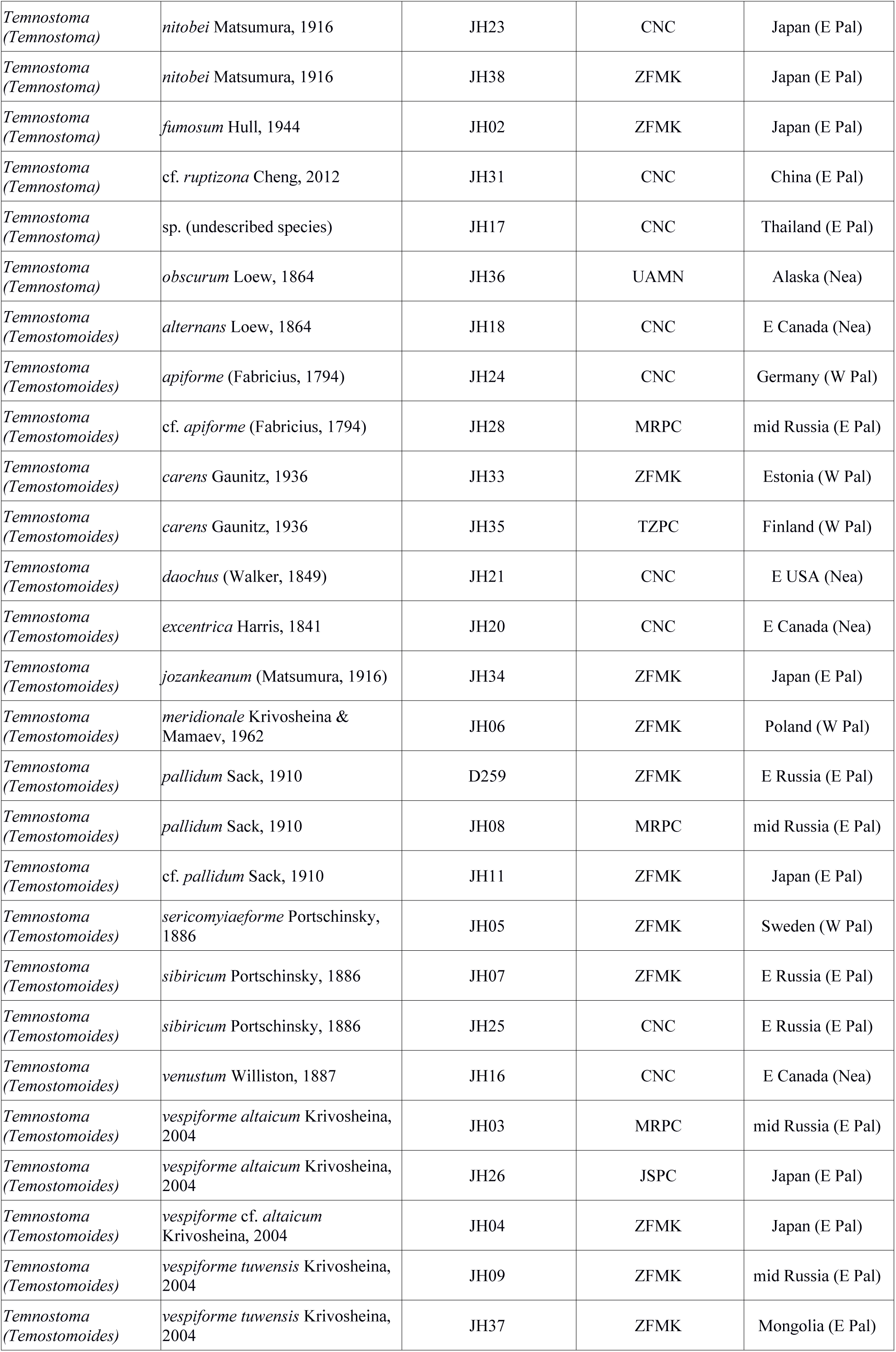
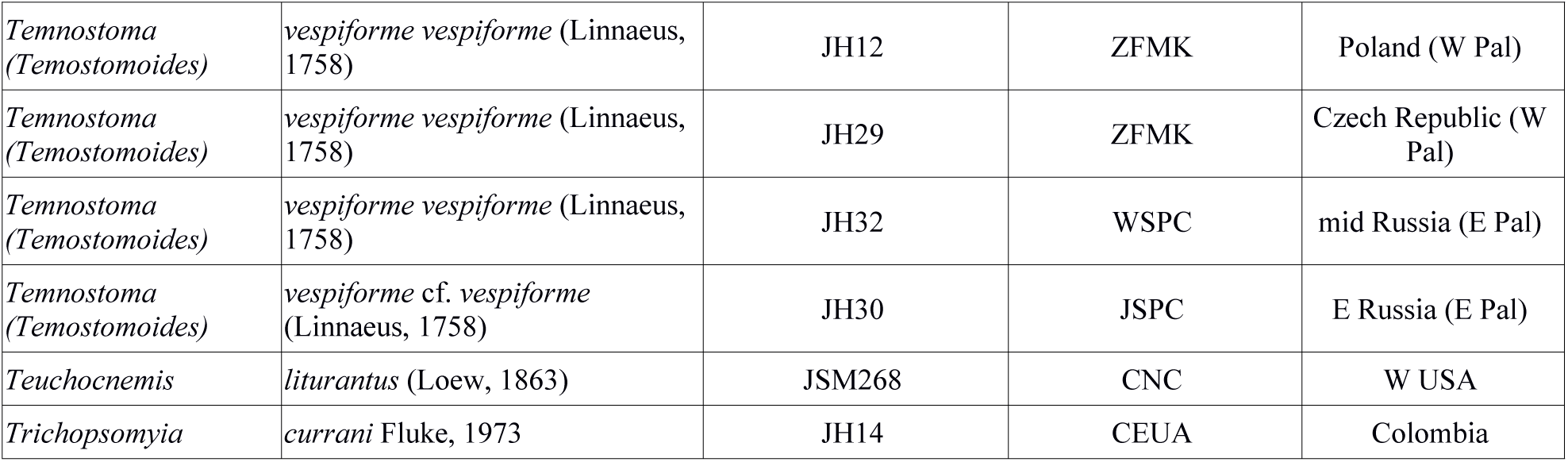
Specimens used in the phylogenetic analysis. For exact locality information and DNA sequence accession numbers, see Supp Table 1.

In our analyses, we included nine species belonging to eight genera other than *Temnostoma*. Two of them are hover flies from different subfamilies: *Trichopsomyia currani* Fluke, 1937 (Pipizinae) and *Allograpta jacqi* Mengual & Ramage, 2018 (Syrphinae), which are resolved sister to each other (Young et al. 2016, Pauli et al. 2018, Moran et al. 2022). These two species were thus used to root the phylogenetic tree. The other outgroup taxa belong to the subfamily Eristalinae. Three of them are hypothesized to be more distant species, namely *Chalcosyrphus chalybeus* (Wiedemann, 1830), *Eristalis arbustorum* (Linnaeus, 1758), and *Spilomyia alcimus* (Walker, 1849). The remaining species are *Takaomyia* sp., *Takaomyia sexmaculata* (Matsumura, 1916), *Teuchocnemis lituratus* (Loew, 1863), and *Pterallastes thoracicus* Loew, 1863; all of them use the front legs to imitate hymenopteran antennae in the same way as *Temnostoma* does and they are traditionally classified into the subtribe Temnostomina (but see results from Moran et al. 2022, who recovered Temnostomina as polyphyletic).

Several different keys were used to identify the specimens. European species were identified using the StN key (Speight and Sarthou 2016) and van Veen (2004). In the case of *Te. bombylans* and *Te. angustistriatum*, high intraspecific variability in characters was found, so the morphology of male genitalia was used to distinguish these species (Krivosheina 2002). For Palaearctic species related to *Te. vespiforme*, the key by Krivosheina (2004) was used. Other species from the Eastern Palaearctic were identified using several keys such as Shiraki (1968), Jilong and Xiping (1995), Huo et al. (2007), Huang and Cheng (2012), and Ichige (2018). For the identification of Nearctic species, Curran (1939), Shannon (1939), and Skevington et al. (2019) were used.

The specimens are held by following institutions: Zoologisches Forschungsmuseum Alexander Koenig, Bonn, Germany (ZFMK), Canadian National Collection of Insects, Arachnids and Nematodes, Ottawa, Canada (CNC), University of Alaska Museum, Fairbanks, USA (UAMN), Universidad de Antioquia, Colombia (CEUA), Menno Reemer personal collection, Leiden, The Netherlands (MRPC), Gerard Pennards personal collection, Leiden, The Netherlands (GPPC), Theo Zeegers personal collection, Leiden, The Netherlands (TZPC), Jeroen van Steenis personal collection, Amersfoort, The Netherlands (JSPC), Wouter van Steenis personal collection, Breukelen, The Netherlands (WSPC).

### 2.2. Genetic data

For the phylogenetic analysis six loci were chosen: the whole mitochondrial COI gene (sequenced in three parts and merged into a single sequence), a part of the large ribosomal unit 28S (D2 region and adjacent parts of D1 and D3 regions) and four nuclear protein-coding genes: Alanyl-tRNA Synthetase (AATS), 5’-end of RNA-binding Protein 15 (RBP-15, also known as “L44”), 5’-end of Casein Kinase 1 (CK1, also known as “L352”), and 5’-end of Tubby-Like Protein (TULP, also known as “L54”). Nuclear protein-coding genes were chosen based on the preliminary results from analyses of Diptera transcriptomes and they were loci with high informative value at the intergeneric level (Young et al. 2016, Pauli et al. 2018, Moran et al. 2022). The list of primers used is available in Table 2.

**Table 2.**
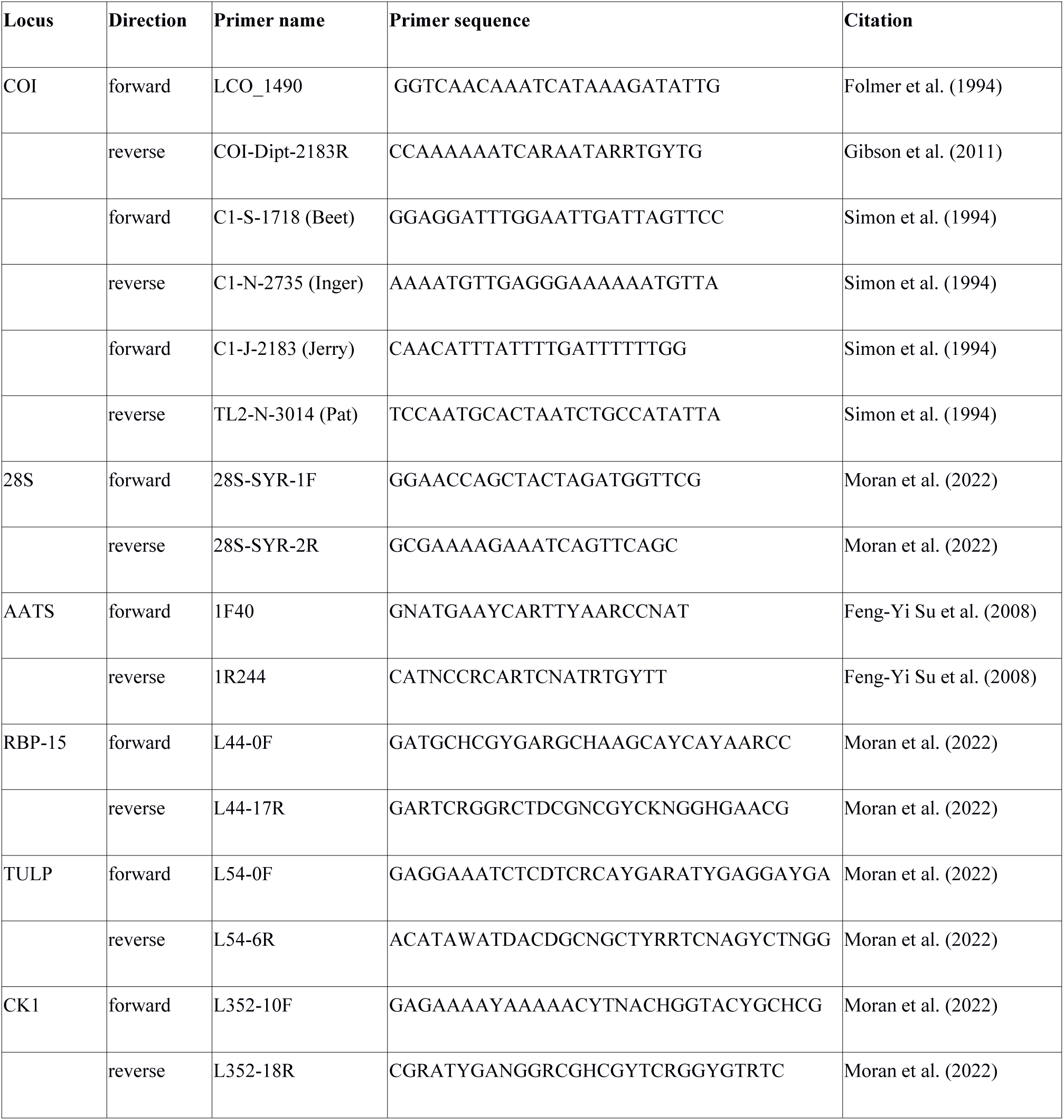
Primers.

The DNA was extracted from a leg from each specimen using the DNeasy Blood & Tissue Kit (QIAgen®) according to the manufacturer’s protocol. Remnants of specimens were preserved and labelled as DNA voucher specimens for the purpose of morphological studies and deposited at the the Canadian National Collection of Insects, Arachnids and Nematodes (CNC) and the Zoological Research Museum Alexander Koenig (ZFMK), as listed in Table 1.

We followed the PCR amplification protocol established by Rozo-Lopez and Mengual (2015). PCR reactions were conducted using Bio-Rad thermal cyclers. The PCR products were sequenced by Macrogen Europe (Amsterdam, The Netherlands) using Sanger sequencing. Sequences were checked for base-call errors, assembled and aligned in Geneious 9.1.7 (Biomatters Ltd.), and all new sequences were submitted to GenBank. see Supp Table 1 for GenBank acession numbers.

Sequences were checked and aligned in Geneious 9.1.7. For protein-coding genes, the MUSCLE alignment algorithm was used in Geneious. In TULP gene, a highly variable intron part was not possible to reliably align and was thus removed prior to the analyses. Hand-made structural alignment was used for 28S (Mengual et al. 2012). Based on the structural alignment of the 28S gene, sequences were divided into two partitions: the first one contains all paired nucleotides (stems) and the second one contains all unpaired nucleotides (loops and connections). The text editor Vim 7.4 was used for partitioning the aligned sequences. For the overview of the partitions used in the analyses, see Table 3.

**Table 3.** The molecular sequences used in MrBayes analysis as separate partitions.

| Partition | Genome | Type | Number of characters (bp) |
| --- | --- | --- | --- |
| COI | mitochondrial | protein-coding | 1420 |
| 28s stems | nuclear | ribosomal | 464 |
| 28s loops | nuclear | ribosomal | 381 |
| AATS | nuclear | protein-coding | 490 |
| CK1 (L44) | nuclear | protein-coding | 1079 |
| TULP (L54) | nuclear | protein-coding | 498 |
| RBP-15 (L352) | nuclear | protein-coding | 671 |
| total |  |  | 5003 |

*2.3. Phylogenetic and statistical analyses*

### 2.3. Phylogenetic and statistical analyses

We conducted Bayesian inference using Markov Chain Monte Carlo (MCMC) in the software MrBayes version 3.2 (Ronquist et al. 2012). To test the robustness of the results, six analyses with different data partitioning strategy and substitutions models were run (Supp Table 3). We used partitioning by gene (as specified in Table 3 with the stems and loops of 28S kept separately, i.e. seven partitions) or by codon (the first and the second codon position combined and the third codon position separated in all protein-coding genes, i.e., 12 partitions). We compared the fit of two substitution models, in both cases with gamma-distributed rate variation across sites and a proportion of invariable sites, the HKY+I+G and GTR+I+G (Arenas 2015), the latter of which often provides the best fit to empirical data (Sumner et al. 2012) by comparing the harmonic mean of the log likelihood and the Akaike information criterion (AIC). We ran the MCMC for 10^8^ steps, extended to 10^9^ steps when the effective sample size was insufficient (<200 for some of the model parameters). We stored the estimated parameter values and a phylogenetic tree every 10,000 steps. For the nexus files with full settings of the analyses, see Supp Files 1-6. The majority rule consensus tree was visualised using FigTree version 1.4.4.

To examine the evolutionary history of mimicry and biogeography, the following characters were obtained for each studied species of *Temnostoma* and *Takaomyia*: biogeography (three levels: Eastern Palaearctic / Western Palaearctic / Nearctic; for the species distributed in more regions, a specimen from each region was included), and morphological characters associated with wasp mimicry, i.e. number of pale stripes per abdominal tergite 2 and 3 (three levels: 1 stripe or less / 2 stripes / 1 normal stripe + 1 dull stripe) and shape of the abdomen (two levels: narrow, i.e. with tergite 2 less than 1.5 times wide as long / wide, i.e. with tergite 2 more than 1.5 times wide as long). The character matrix is available in Supp Table 2. These characters were mapped onto the phylogenetic tree in software R version 4.2.0 (R Core Team 2022) using the package ‘phytools’ (Revell 2012) for the mapping of morphological traits and the package ‘BioGeoBEARS’ (Matzke 2018) for the analyses of biogeography following the example script provided by Matzke (2018). Prior to these analyses, we pruned the tree to exclude taxa other than *Temnostoma* and *Takaomyia* (in order to prevent distant species and sparsely sampled parts of the Temnostomina tree from affecting the results) and to keep only one specimen per species (or putative species) or to keep one specimen from each biogeographical region in the case of widely distributed species. We first tested for the presence of a phylogenetic signal in the morphological traits using the measure of phylogenetic signal in binary traits, D (Fritz and Purvis 2010), implemented in the function phylo.d() in the R package ‘caper’ (Orme et al. 2018). We then reconstructed the ancestral states of both morphological traits using the function ace() in phytools and tested whether the evolution of body shape and colouration is correlated using the Pagel’s test implemented in the function fitPagel() in phytools. To reconstruct the biogeography of *Temnostoma* and *Takaomyia*, we compared the fit of six biogeographical models implemented in BioGeoBEARS using the Akaike information criterion (AICc) (Akaike 1973): the dispersal–extinction–cladogenesis model (DEC) (Ree and Smith 2008), a model based on the dispersal-vicariance model of Ronquist (1997) (“DIVA-like”) and a model based on the BayArea model of Landis et al. (2013) (“BayArea-like”). We also tested a modified version of each of these three models to include the parameter J, which adds the process of jump dispersal at speciation (Matzke 2014, Matzke 2021).

Focus stacked images were created using the software Zerene Stacker ® ver. 1.04 (Richland, Washington, USA), based on photographs of selected pinned specimens taken with a Canon EOS 7D® mounted on a P–51 Cam-Lift (Dun Inc., VA, USA) and with the help of Adobe Lightroom ® ver. 5.6.

## 3. Results

### 3.1. Phylogeny

We obtained a Bayesian phylogenetic tree with highly supported topology. The model with the highest log likelihood and the lowest AIC (Supp Table 3) had partitioning by codon, GTR substitution model with gamma-distributed rate variation across sites and a proportion of invariable sites (“GTR + I + Γ” or “GTR + I + G” model) and was used in the results reported in the main text. The consensus trees obtained using all combinations of the model settings are available in Supp Files 7-12. The final 50% majority rule consensus tree is displayed in Fig. 1. The topology of the tree also remains stable with different settings of the phylogenetic analyses (Supp Table 3), such as data partitioning by gene or codon and the substitution model used (Supplementary Files 7-12).

**Figure 1.**
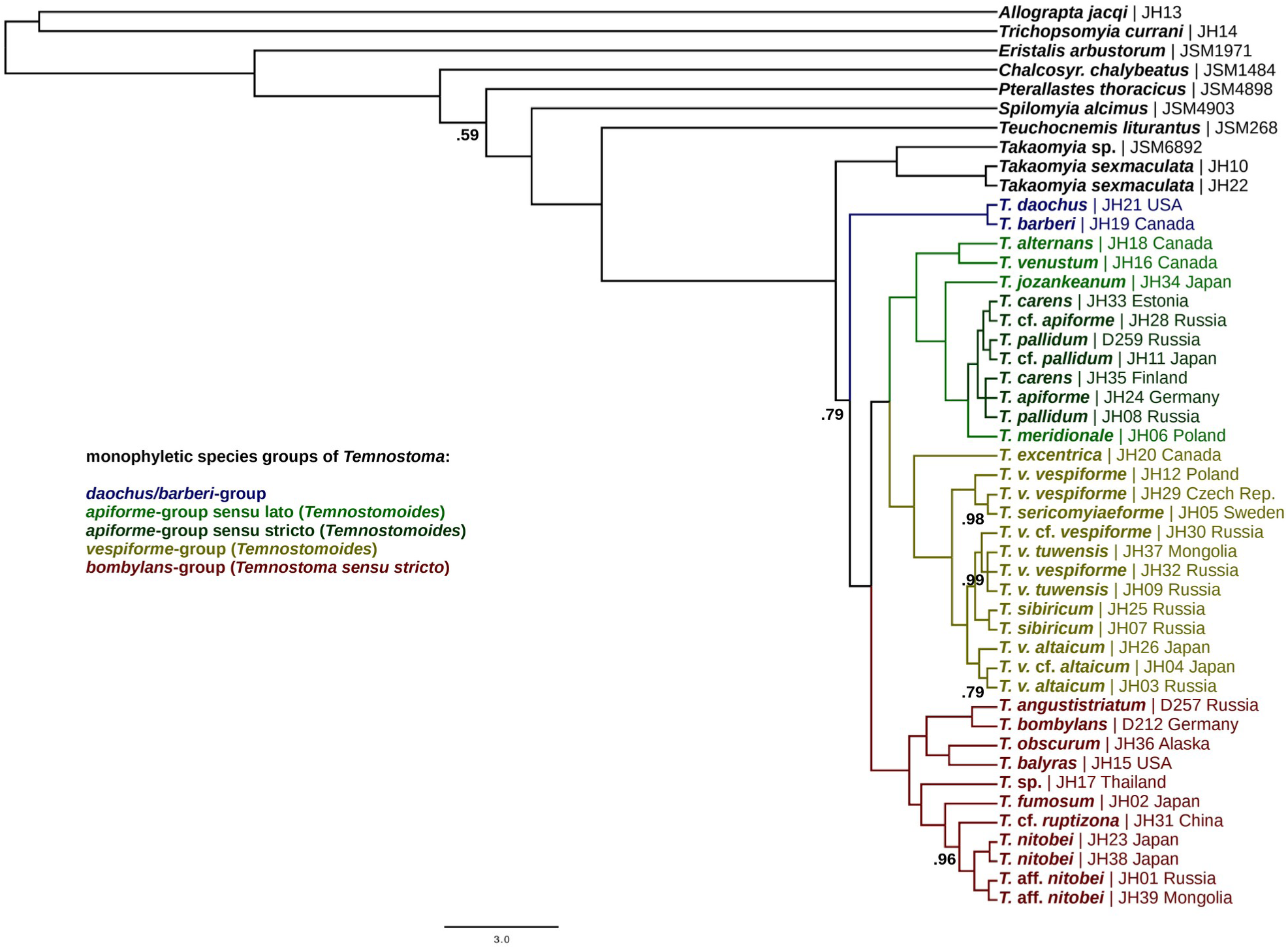
Majority-rule consensus phylogenetic tree from the multigene Bayesian analysis. In taxon names, “*T.*” stands for *Temnostoma* and “*v.*” stands for *vespiforme*. Species groups are highlighted by various colours. Posterior probabilities of most stems are equal to 1, exceptions are marked in the tree.

Our results support the monophyly of genera *Temnostoma* and *Takaomyia*, which are sister to each other. The remaining genera of Temostomina (*Teuchocnemis lituratus* and *Pterallastes thoracicus*) are not closely related to this clade. Except for two Nearctic species, *Temnostoma daochus* and *Te. barberi*, which are sister to each other and together form a clade sister to the rest of the genus *Temnostoma*, both subgenera and all species groups were found to be monophyletic. Several species of *Temnostoma (Temnostomoides)* that have not been assigned to any species group yet (i.e. *Te. jozankeanum*, *Te. venustum*, and *Te. alternans*), as well as *Te. meridionale*, turned out to be closely related to the *apiforme*-group, and we thus call them “*apiforme*-group *sensu lato*”.

### 3.2. Evolution of morphological characters

Across species, there is a strong phylogenetic signal in the distribution of morphological characters connected to mimicry, i.e. the shape of the abdomen and the number of stripes per abdominal tergite. Body shape (narrow or wide abdomen) had the measure of phylogenetic signal in binary traits, D = -1.50 (P<0.001). The number of stripes on abdominal tergites (one or two) had the value of D = -1.10 (P<0.001). Moreover, both characters were highly correlated based on the Pagel’s binary character correlation test (dependent x independent, likelihood ratio = 19.50, P = 0.0006), so the species are usually either good wasp mimics (having wide abdomen as well as two yellow stripes per tergite), or imperfect mimics (having narrow abdomen as well as dark colour pattern). There are a few exceptions, which have wide abdomens, but only a single stripe (*Temnostoma sericomyiaeforme*) or two stripes, but the second one reduced into a narrow dull band (*apiforme*-group *sensu stricto*). In all these cases, secondary reduction of the colouration appeared to be the most plausible explanation of the pattern. The phylogenetic tree with estimated character states in ancestral nodes is shown in Fig. 2.

**Figure 2.**
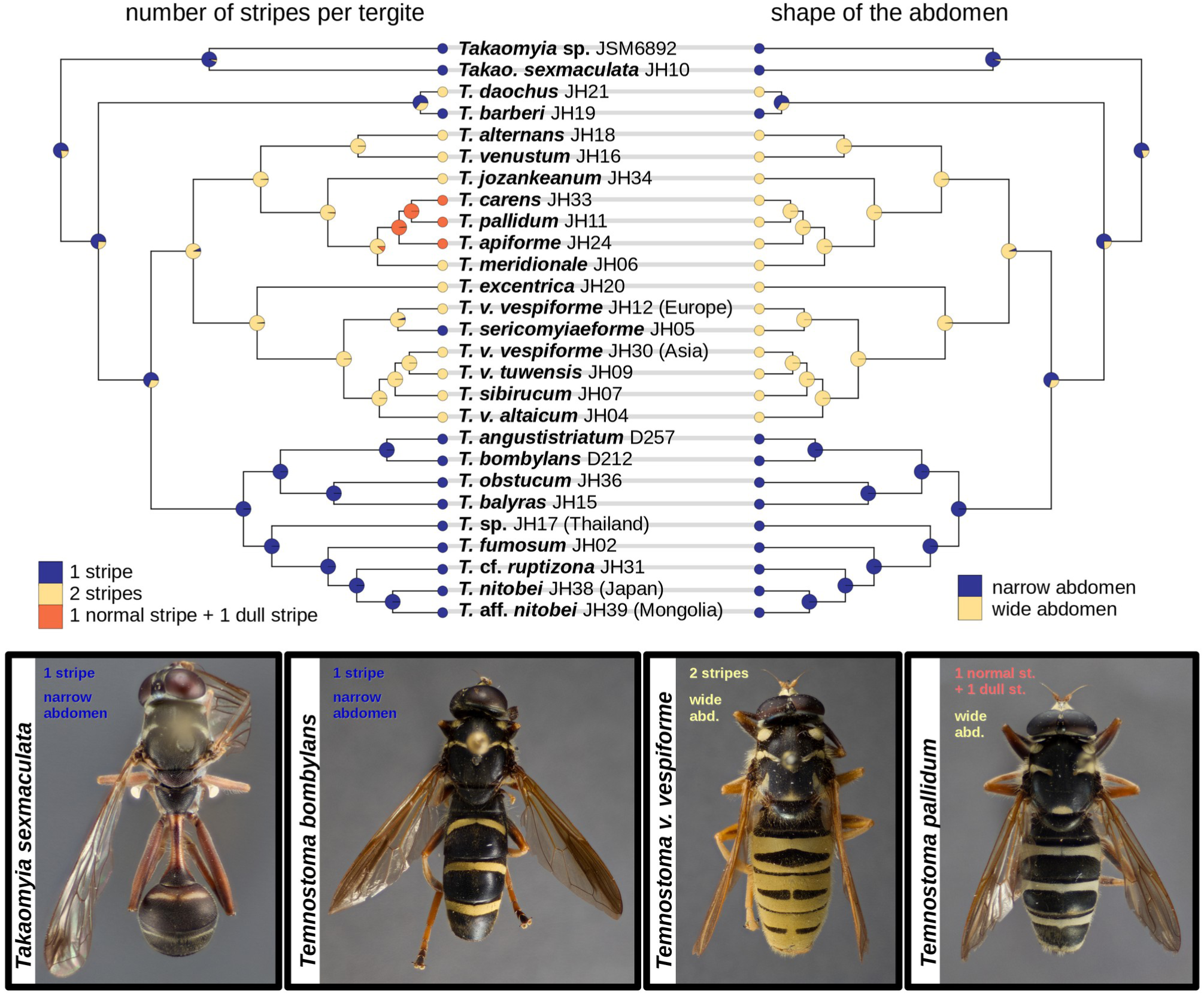
Evolution of the morphological characters in *Temnostoma* and *Takaomyia*. Only one specimen per species or a few specimens from various geographic areas are used in the tree.

### 3.3. Biogeography

The best biogeographical model according to AICc was the DIVA-like+J model, but DEC+J and BayArea- like+J had only marginally higher AICc (ΔAICc<2, Table 4). Ancestral areas estimated by these three models were also very similar (Fig. 3 and Supp Figs 1-2). The region of origin of the last common ancestor of the *Takaomyia* and *Temnostoma* clade is unclear, the highest probability is assigned to a wide distribution in the Eastern Palaearctic and the Nearctic, but the last common ancestor of the genus *Temnostoma* most likely inhabited the Nearctic Region (Fig. 3). Within each of the three main species groups, there is always one deep divergence dividing a Nearctic and a Palaearctic lineage, representing three distinct dispersal events from the Nearctic to the Eastern Palaearctic taking place during a similar time period and the results also suggest one more recent dispersal from the Eastern Palaearctic back to the Nearctic (Fig. 3). Dating of these events is, however, not available.

**Figure 3.**
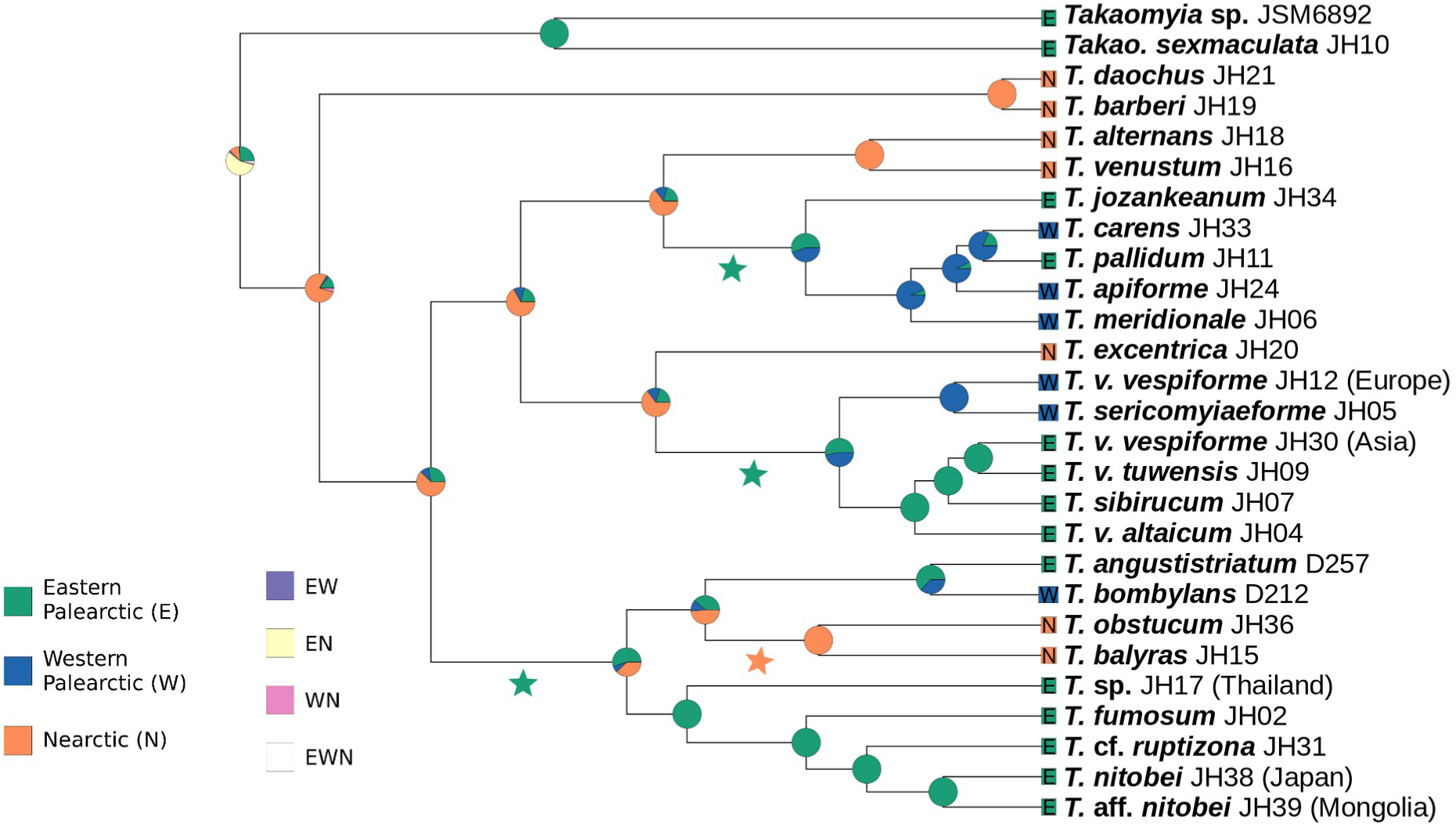
Biogeography of *Temnostoma* and *Takaomyia*. Only one specimen per species or a few specimens from various geographic areas are used in the tree. BioGeoBEARS model DIVALIKE+J, ancestates: global optim, 3 areas max. d=0, e=0, j=0.1303, LnL=-23.69. Inferred dispersal events are marked by stars (green for dispersal to the Eastern Palaearctic and orange for dispersal to the Nearctic).

**Table 4.** Comparison of six models of the biogeographical reconstruction computed by the BioGeoBEARS software: dispersal–extinction–cladogenesis model (DEC), dispersal-vicariance analysis (DIVA-like) and BayArea-like model and their modified versions, including the parameter J, which represents per-event weight of founder events. The model with the lowest AICc is highlighted in bold. Parameters of the models are d (dispersal) describing the rate of range expansion, e (extinction) mirroring potential range contraction and j, which represents founder effects.

| Model | log likelihood | No. parameters | d | e | j | AICc | ΔAICc |
| --- | --- | --- | --- | --- | --- | --- | --- |
| **DIVA-like+J** | **-23.69** | **3** | **1\*10<sup>-12</sup>** | **1\*10<sup>-12</sup>** | **0.13** | **53.38** | **0** |
| DEC+J | -23.96 | 3 | 1*10<sup>-12</sup> | 1*10<sup>-12</sup> | 0.13 | 53.92 | 0.54 |
| BayArea-like+J | -24.66 | 3 | 1*10<sup>-7</sup> | 1*10<sup>-17</sup> | 0.13 | 55.32 | 1.94 |
| DIVA-like | -33.3 | 2 | 0.09 | 1*10<sup>-12</sup> | 0 | 70.6 | 17.22 |
| DEC | -36.75 | 2 | 0.07 | 1*10<sup>-12</sup> | 0 | 77.51 | 24.13 |
| BayArea-like | -47.24 | 2 | 0.06 | 0.34 | 0 | 98.48 | 45.1 |

## 4. Discussion

### 4.1. Phylogeny

*Temnostoma* and *Takaomyia* were found to be two monophyletic genera sister to each other, but unrelated to other genera of the subtribe Temnostomina, which is consistent with Moran et al. (2022). The monophyly of both subgenera and all three species groups of *Temnostoma* were supported, with the notable exception of two North American species (i.e., *Temnostoma daochus* and *Te. barberi*), which were found to be closely related to each other and forming a lineage sister to the rest of the genus. Bellow, we first discuss the implications of our results for our understanding of the evolution of mimicry in *Temnostoma* and its biogeography, and later turn out attention to issues of the systematics of *Temnostoma*.

### 4.2. Evolution of mimicry

Based on the estimation of ancestral morphological characters related to mimicry, we suggest that the last common ancestors of *Temnostoma* + *Takaomyia* had narrow abdomens and dark colour patterns, and were thus either imperfect mimics, or mimics of Hymenoptera with similar appearances (such as Ropalidiini or Eumeninae). The perfect mimicry of social wasps of the subfamily Vespinae evolved later, and two times independently within the genus *Temnostoma*: for the first time, it appeared on the stem lineage of *Temnostoma (Temnostomoides)*, which then dispersed across the North hemisphere, and for the second time, it appeared very recently in a North American species, *Te. daochus*. Leavey et al. (2021) estimated the dynamics of perfect wasp mimicry evolution in hover flies and they concluded that perfect mimicry appeared several times independently in hover flies. Here, our results suggest that the mimicry evolved even faster in hover flies as perfect wasp mimicry appeared two times independently within a single and relatively species-poor genus. The evolutionary rates of changes in mimicry-related morphological characters seem to vary a lot across the phylogenetic tree. For instance, while in *daochus/barberi* group (Fig. 4), there is a mimicry switch on the level of two genetically almost identical species, in the other species groups, the morphology is highly conserved. Moreover, in *Te. daochus*, there is also sexual dimorphism in the shape of abdomen (abdomen of male is wider than abdomen of female).

**Figure 4.**
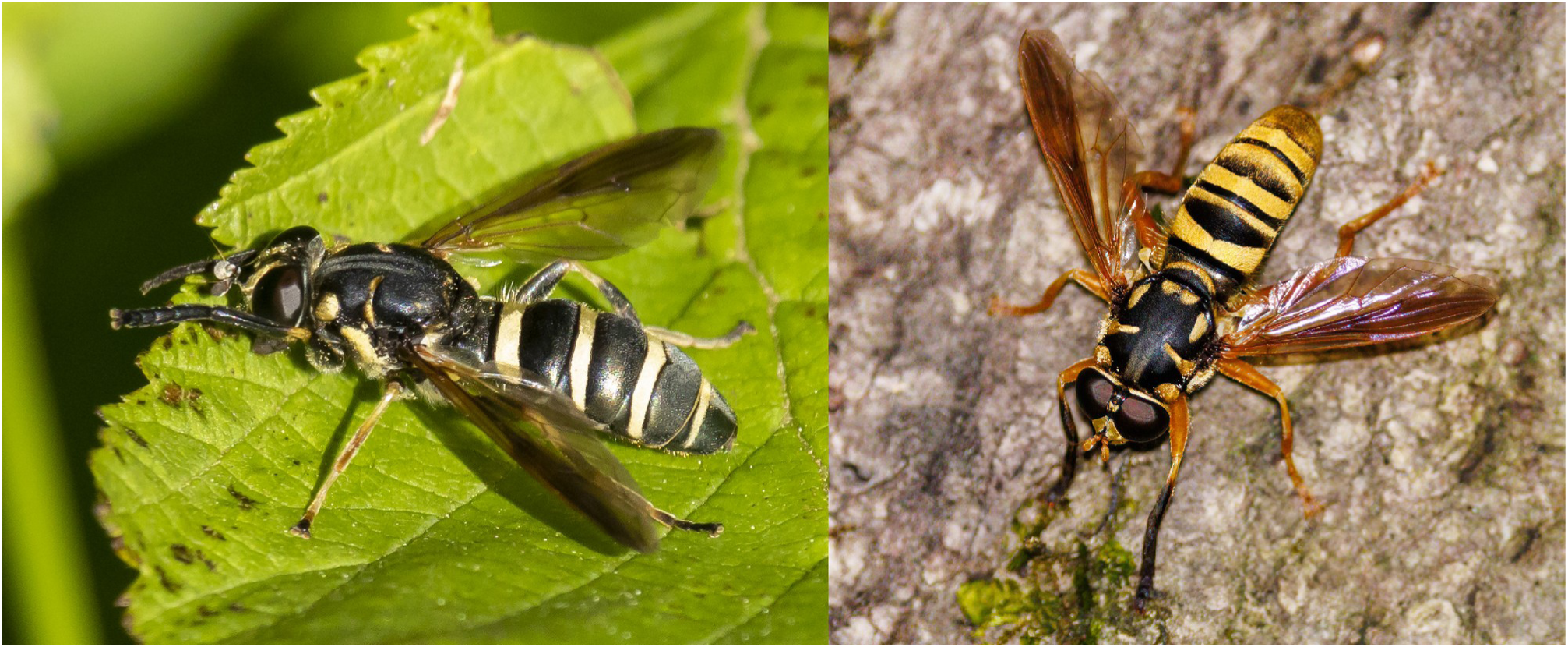
*Temnostoma barberi* (left, photo by Denis Doucet) and *Te. daochus* (right, photo by Brad Moon): two closely related Nearctic species exhibiting striking difference in their mimicry.

The majority of *Temnostoma* species exhibiting perfect wasp mimicry are assumed to imitate yellowjackets (genera *Vespula* Thomson, 1869 and *Dolichovespula* Rohwer, 1916). An exception occurs in Japanese species *Temnostoma jozankeanum* (*apiforme*-group *sensu latu*). Even though the mapped morphological characters remain in the *Temnostomoides* plesiomorphic state in this species (i.e. two stripes per tergite and wide abdomen), *Te. jozankeanum* differs from the others by higher proportion of yellow colouration and larger size, so its general appearance is more striking compared to other *Temnostoma* species and it resembles hornets (genus *Vespa* Linnaeus, 1759).

Secondary reduction of the perfect wasp mimicry appeared two times independently within *Te. (Temnostomoides)*, specifically in *Te. sericomyiaeforme* and in lesser degree in the *apiforme*-group *sensu stricto*. As both taxa have boreal distribution, our findings are in congruence with Taylor’s hypothesis that the dark pattern could emerge as trade-off between mimicry and thermoregulation (Taylor et al. 2016). Consistently, also in the *daochus/barberi* group, the brighter and more convincing mimetic species *Te. daochus* has its range shifted more to the south compared to the darker *Te. barberi* (Skevington et al. 2019). A higher proportion of melanic individuals in higher latitudes are documented in many other species of hover flies such as *Melanostoma* (Haarto and Ståhls 2014).

We confirmed that the behavioural mimicry (imitation of Hymenopteran antennae with front legs) is an ancestral state that all species of *Temnostoma* inherited from a common ancestor of *Temnostoma* + *Takaomyia* and it remained unchanged across several evolutionary changes in morphology, which led to switches in mimicry models and mimicry perfectness. We can clearly conclude that the behavioural mimicry must have preceded the appearance of perfect wasp mimicry in *Temnostoma* and is more evolutionary conservative than the morphological characters. Consistently with Moran et al. (2022), our results suggest that the behavioural mimicry in *Temnostoma* + *Takaomyia* evolved independently of similar behaviour known in other Temnostomina species such as *Teuchocnemis lituratus* and *Pterallastes thoracicus*, and in *Spilomyia*, where intrageneric variability in behavioural mimicry is known (Penney et al. 2014).

### 4.3. Biogeography

All the three main *Temnostoma* species groups (*bombylans*-group, *vespiforme*-group, and *apiforme*-group *sensu lato*) have their representatives in the Western and the Eastern Palaearctic as well as in the Nearctic. One dispersal event between an American and a Eurasian lineage was found within each of the three species groups, and with the exception of the boreal *apiforme*-group *sensu stricto*, one dispersal event between the Eastern and the Western Palaearctic was found in each species group. It seems that the formation of *vespiforme*-like (2 stripes per tergite and wide abdomen) and *bombylans*-like (1 stripe per tergite and narrow abdomen) morphotypes took place before the morphotypes dispersed across Eastern and Western hemispheres, but *apiforme*-like (1 normal stripe + 1 dull stripe, wide abdomen) evolved in situ.

Interestingly, all three divergences between the American and Eurasian lineages within each species group are estimated to take place during a similar time period. Unfortunately, our current data do not allow us to make a reliable estimation of the divergence age, because of the lack of fossil record or reliably dated higher phylogeny of the family Syrphidae. Even though Statz (1940) described a fossil hover fly *Temnostoma sacki* from deposits of late Oligocene, we have doubts about the classification of this species into the genus, in particular due to enlarged femora, which are not present in any recent species of *Temnostoma* or *Takaomyia*. Consequently, there is no fossil record that could be used as a calibration point in our tree. We assume that the dispersal between the Eastern Palaearctic and the Nearctic regions might have happened during the presence of the second (from the Miocene to the late Pliocene, 14–3.5 Ma) or the third (the Pleistocene, 1.5–1 Ma) Beringian Land Bridge (Sanmartín et al. 2001), similarly to other insects with comparable geographic distribution, such as various butterfly genera (e.g., Peña et al. 2015, Kleckova et al. 2015).

### 4.4. Systematics

In the *bombylans*-group, contrary to the other species groups, the deepest divergence was found between the East Asian lineage and the European+American lineage. However, better taxon sampling would be needed to fully understand the deep disjunction between species of the Western and Eastern Palaearctic. Moreover, *Te. angustistriatum*, which is morphologically very similar to and has sympatric distribution with *Te. bombylans* (Krivosheina and Ståhls 2003), appeared to be genetically very closely related to *Te. bombylans*. Based on additional COI sequences of both species from the BOLD database (https://boldsystems.org), there is no clear molecular disjunction between the two species, and we found several specimens with intermediate morphological characters. More detailed taxonomical revision will be needed, but the current evidence suggests that these taxa might be a single species with high morphological variability.

Consistently with Krivosheina (2004), the *vespiforme*-group consists of *Te. vespiforme vespiforme*, *Te. v. altaicum*, *Te. v. tuwensis*, *Te. sericomyiaeforme*, and *Te. sibiricum*. Moreover, one Nearctic species was found to be closely related to this group, *Temnostoma excentrica*. This species is sometimes called in literature by its junior synonym name *Te. aequalis* Loew, 1864, or incorrectly identified as the Palaearctic species *Te. vespiforme* (Krivosheina 2012). Moreover, even the Palaearctic *Te. vespiforme* and even its nominative subspecies *Te. v. vespiforme* appears to be a paraphyletic taxon, which includes *Te. sericomyiaeforme* and *Te. sibiricum*. Based on our results, the phylogenetic relations within this group reflect geographical distribution rather than morphological characters: European *Te. v. vespiforme* seems to be the same taxon as *Te. sericomyiaeforme* and the Asian *Te. v. vespiforme* (as defined by Krivosheina 2004) is the same taxon as *Te. v. tuwensis*. Moreover, we did not find any support for treating some taxa of the *vespiforme*-group as subspecies and other as species. Based on our inferred phylogeny, *Te. v. tuwensis* (including Asian *Te. v. vespiforme*) and *Te. v. altaicum* should be treated as separate species as they are more closely related to *Te. sibiricum* than Linnaeus’s *Te. vespiforme*. On the other hand, *Te. sericomyiaeforme* is an inner group of European *Te. v. vespiforme* and should not be thus treat as separate species, but rather as a Nordic variety of the European *Te. v. vespiforme*. Treating *Te. sericomyiaeforme* as junior synonym of *Te. vespiforme* is also supported by several records of mating behaviour between *Te. sericomyiaeforme* and *Te. vespiforme* (e.g. https://inaturalist.laji.fi/observations/19048822 and https://www.inaturalist.org/observations/36833709). Alternatively, *Te. sibiricum* together with *Te. sericomyiaeforme* could both be treated as junior synonyms of *Te. vespiforme*, and so the *vespiforme*-group would consist of two species only (*Te. vespiforme* and *Te. excentrica*).

The *apiforme*-group *sensu stricto* (i.e., the *apiforme*-group as defined by Krivosheina 2004) was found to be monophyletic. Within this group, three species have been distinguished based on morphology (Krivosheina 2003, 2004): *Te. apiforme*, *Te. carens*, and *Te. pallidum*. However, the genetic data and geographical patterns do not support these three taxa. This lineage rather looks like a genetically uniform and morphologically variable taxon with Boreal distribution. Consistently with morphological studies by Krivosheina (2005), *Te. meridionale*, which closely resembles *Te. vespiforme* by the outer morphology, but it is similar to the *apiforme*-group based on male genitalia, appeared to be the sister species to the *apiforme*-group *sensu stricto*. Such phylogenetic placement may indicate that the reduction of the posterior yellow stripe on abdominal tergites and consequent reduction of the perfectness of the mimicry is secondary in this lineage. This hypothesis is also supported by close relativeness with other perfect mimics, such as the East Asian species *Te. jozankeanum* resembling hornets of the genus *Vespa* Linnaeus, 1758, and the North American yellow-jacket perfect mimics *Te. alternans* and *Te. venustum*.

The most surprising result is the close relation between two American species *Te. barberi* and *Te. daochus*. *Temnostoma barberi* is a dark, narrow species previously classified in the *bombylans*-group (Curran 1939, Shannon 1939), while *Te. daochus* is a remarked wasp mimetic with wide abdomen and with an even brighter colour pattern than most of the representatives of the *vespiforme*-group and the *apiforme*-group, and it was thus classified into the subgenus *Te. (Temnostomoides)*. Furthermore, the genetic distance between the two taxa is extraordinarily low; there are only very few substitutions in the studied sequences, similar to intraspecific variability in other two-taxa comparisons. The results were double-checked for a mistake, such as contamination or lab-notes confusion, but no such error was revealed and multiple DNA barcodes support this discovery. Moreover, we found several morphological characters supporting the monophyly of this clade (absence of pilosity on the metasternum, structure of the pilosity on the face, wing dark marking common in both species). Such findings suggest a very fast evolutionary shift between perfect and imperfect mimicry, and the evolution of this lineage should be studied more in detail as a model for microevolutionary changes in mimicry perfectness.

To summarise, our molecular phylogeny illustrated several taxonomic issues in *Temnostoma*. However, we do not want to propose any formal taxonomical changes in this paper. A thorough taxonomical revision, including more detailed morphological study and barcode data of a higher number of specimens from problematic taxa will be needed.

## 5. Conclusions

Our results show that the evolution of morphological traits related to mimicry is highly dynamic in hover flies, as two independent origins of perfect wasp mimicry and two independent reductions of mimicry accuracy have been documented within a single genus with relatively low number of species. Even though each of the morphotypes is widely distributed across the Holarctic, each of the main lineages underwent only one dispersal event between the Nearctic and Palaearctic Realms.

## CRediT authorship contribution statement

Jiří Hadrava: Conceptualization, Methodology, Software, Validation, Formal analysis, Investigation, Visualization, Writing - Original Draft, **Jan Klečka**: Methodology, Software, Formal analysis, Visualization, Writing - Review & Editing, Supervision, **Kevin Moran:** Methodology, Validation, Writing - Review & Editing, **Irena Klečková:** Software, Formal analysis, **Scott Kelso:** Methodology, Investigation, **Claudia Etzbauer:** Methodology, Investigation, **Jeffrey H. Skevington:** Methodology, Validation, Investigation, Writing - Review & Editing, **Ximo Mengual:** Conceptualization, Methodology, Validation, Investigation, Writing - Review & Editing, Supervision

## Acknowledgements

We thank Martin Hauser, Katsuyoshi Ichige, Lukasz Mielczarek, Augusto L. Montoya, Gerard Pennards, Menno Reemer, Derek Sikes, Fredrik Sjöberg, Jozef Slowik, John T. Smit, Gunilla Ståhls, Axel Ssymank, André van Eck, Jeroen van Steenis, Wouter van Steenis, and Theo Zeegers for providing the specimens. The photographs were provided by Denis Doucet and Brad Moon. We thank to Björn Rulik for comments and suggestions. The work was supported by German Academic Exchange Service (DAAD), Czech Science Foundation (GA20-14872S), Institutional Research Support grant of the Charles University, Prague (SVV260571/2022), and Charles University Research Centre program No. 204069.

## Supplementary Data

Supplementary data are available online.

