## Supplementary material for "The evolution of wasp mimicry and biogeography in the genus *Temnostoma* (Diptera: Syrphidae)": Hadrava_etal_2024_Supplementary_material.pdf

**Supplementary Table 1 –** List of all specimens used in our analyses with taxonomic information, GenBank accession numbers of all DNA sequences, hosting institution with specimen ID, and detailed information on collection sites. The table is in a separate file.

**Supplementary Table 2 –** Traits and biogeographic distribution of individual species. The table is in a separate file.

**Supplementary Table 3 –** A comparison of six MrBayes models with different settings used in our analyses. nst = substitution model setting in MrBayes (nst=2 is the HKY model, nst=6 is the GTR model, and nst=mixed allows either model jumping, i.e., the MCMC sampler explores different models and weights the results according to the posterior probability of each model; see Ronquist et al. 2012). Gamma-distributed rate variation across sites and a proportion of invariable sites was implemented in all models. Model 5, with partitioning by codon, GTR substitution model (nst=6) with gamma-distributed rate variation across sites and a proportion of invariable sites ("GTR + I + Γ" or "GTR + I + G" model), had the highest log likelihood value (in bold) and was used in the results reported in the main text. The consensus trees obtained using all combinations of the model settings are shown in Figures S1-S6.

| model | partitioning | substitution model | nst | log likelihood (harmonic mean) | no. parameters | AIC | ΔAIC |
|---|---|---|---|---|---|---|---|
| model 1 | by gene | HKY+G+I | 2 | -24820.90 | 59 | 49760 | -3136 |
| model 2 | by gene | GTR+G+I | 6 | -24612.50 | 100 | 49425 | -2800 |
| model 3 | by gene | mixed | mixed | -24641.49 | 107 | 49497 | -2873 |
| model 4 | by codon | HKY+G+I | 2 | -23440.24 | 99 | 47078 | -454 |
| **model 5** | **by codon** | GTR+G+I | **6** | **-23142.01** | **170** | **46624** | **0** |
| model 6 | by codon | mixed | mixed | -23160.36 | 182 | 46685 | -61 |

**The following files contain the nexus files for individual models listed in Supplementary Table 3:**

**Supplementary File 1:** Temnostoma_MrBayes_model_1.nex
**Supplementary File 2:** Temnostoma_MrBayes_model_2.nex
**Supplementary File 3:** Temnostoma_MrBayes_model_3.nex
**Supplementary File 4:** Temnostoma_MrBayes_model_4.nex
**Supplementary File 5:** Temnostoma_MrBayes_model_5_BEST_MODEL.nex
**Supplementary File 6:** Temnostoma_MrBayes_model_6.nex

**The following files contain the consensus trees for individual models listed in Supplementary Table 3:**

**Supplementary File 7:** Temnostoma_MrBayes_model_1.nex.con.tre
**Supplementary File 8:** Temnostoma_MrBayes_model_2.nex.con.tre
**Supplementary File 9:** Temnostoma_MrBayes_model_3.nex.con.tre
**Supplementary File 10:** Temnostoma_MrBayes_model_4.nex.con.tre
**Supplementary File 11:** Temnostoma_MrBayes_model_5_BEST_MODEL.nex.con.tre
**Supplementary File 12:** Temnostoma_MrBayes_model_6.nex.con.tre

**Supplementary Figure 1 –** Biogeography of *Temnostoma* and *Takaomyia* (DEC+J model).

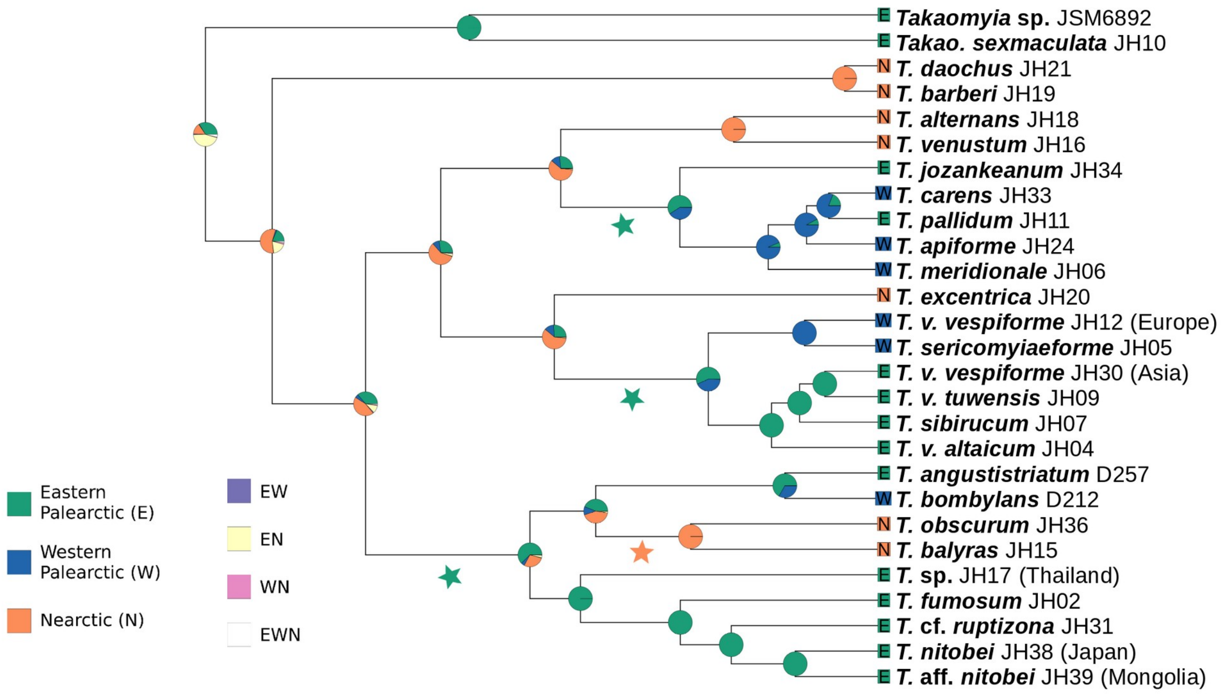


**Supplementary Figure 2 –** Biogeography of *Temnostoma* and *Takaomyia* (BAYAREALIKE+J model).

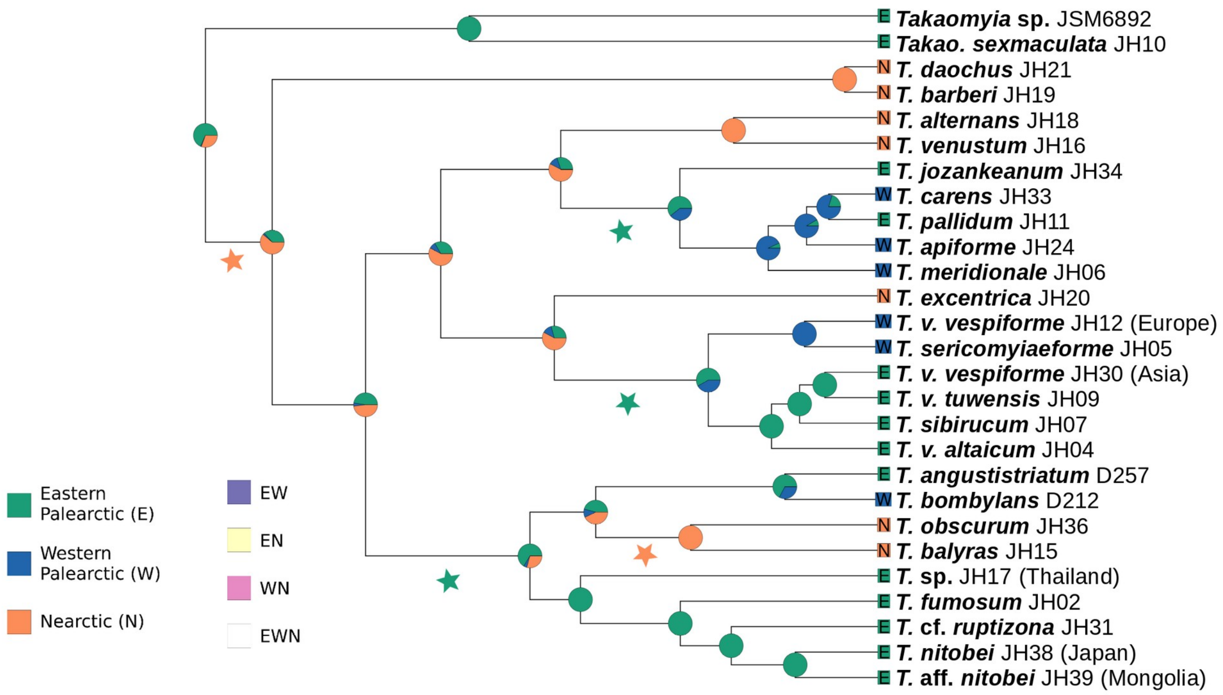
